# The Integration of Proteogenomics and Ribosome Profiling Circumvents Key Limitations to Increase the Coverage and Confidence of Novel Microproteins

**DOI:** 10.1101/2023.09.27.559809

**Authors:** Eduardo V. de Souza, Angie L. Bookout, Christopher A. Barnes, Brendan Miller, Pablo Machado, Luiz A. Basso, Cristiano V. Bizarro, Alan Saghatelian

## Abstract

There has been a dramatic increase in the identification of non-conical translation and a significant expansion of the protein-coding genome and proteome. Among the strategies used to identify novel small ORFs (smORFs), Ribosome profiling (Ribo-Seq) is the gold standard for the annotation of novel coding sequences by reporting on smORF translation. In Ribo-Seq, ribosome-protected footprints (RPFs) that map to multiple sites in the genome are computationally removed since they cannot unambiguously be assigned to a specific genomic location, or to a specific transcript in the case of multiple isoforms. Furthermore, RPFs necessarily result in short (25-34 nucleotides) reads, increasing the chance of ambiguous and multi-mapping alignments, such that smORFs that reside in these regions cannot be identified by Ribo-Seq. Here, we show that the inclusion of proteogenomics to create a Ribosome Profiling and Proteogenomics Pipeline (RP3) bypasses this limitation to identify a group of microprotein-encoding smORFs that are missed by current Ribo-Seq pipelines. Moreover, we show that the microproteins identified by RP3 have different sequence compositions from the ones identified by Ribo-Seq-only pipelines, which can affect proteomics identification. In aggregate, the development of RP3 maximizes the detection and confidence of protein-encoding smORFs and microproteins.

## Introduction

Small Open Reading Frames (smORFs, <300-500 nucleotides)^1–4^ were excluded from typical genome annotation workflows because there were no methods to validate their translation, which led to the concern that most smORFs were false positives^5,6^. The advent of Ribosome profiling, or Ribo-Seq, provided empirical data to identify actively translated smORFs and provide an accurate snapshot of translation at sub codon resolution^7,8^. The ability to accurately define translated smORFs has led to greater and greater efforts in this area to identify and functionally characterize these genes.

The assignment of smORFs occurs after Ribo-Seq data is fed into a bioinformatics workflow. A variety of different scoring algorithms are used to identify smORFs, such as RibORF^9^, Ribocode^10^, RiboDIPA^11^, ORFRater^12^, RiboTaper^13^, ORFscore^14^, and RP-BP^15^. These tools differ substantially in their approach, but most tools were designed for the *de novo* annotation of the translatome by assessing the 3-nucleotide periodicity, although machine learning algorithms trained on available annotation data are also common. The performance of these tools vary in many metrics, and ORFRater performance is also heavily affected by the usage of the drug harringtonine during sample preparation, which is a protein synthesis inhibitor that stalls the ribosome on the initiation codon to better identify the start of a (sm)ORF^12^.

Although many pipelines are being successfully used for the identification of actively translated smORFs, there are some shortcomings that limit smORF annotation. A common problem is the existence of reads that map to multiple locations in the genome during the alignment step of these workflows^16^. These duplicated regions might result from a variety of events, such as whole genome duplication^17,18^, recombination^19^, retro-transposition^20^ and gene duplication^21^. Multi-mapping reads must be addressed with caution during genomics and transcriptomics analysis, as they can confound gene quantification and make genome annotation more difficult.

In a typical RNA-Seq analysis with short-reads, multi-mapping reads make up from 5 to 40% of total mapped reads ^16^. This proportion is expected to be bigger in a Ribo-Seq analysis, as reads from ribosome footprints range from 25 to 34 bp^2,7^, which increases the chance of them mapping to multiple sites when compared to traditional Illumina short-reads, which are 75 to 300 bp long^22^. Most of the time, multi-mapped reads are discarded during read assignment^16^, which limits the number of novel smORFs that can be found to be actively translated, especially when working with a genome with a high number of paralogous sequences.

Proteogenomics is a multi-omics approach that integrates genomics, transcriptomics and proteomics, and can be used for a multitude of tasks, including supporting genome annotation and allowing the identification of microproteins encoded by smORFs^23,24^. The most robust evidence for translation of a smORF is the direct detection of the resultant microprotein. As a consequence, while the total number of smORFs revealed by proteogenomics is eclipsed by the higher number of Ribo-Seq annotated smORFs, there is added value in validating the microprotein translation with proteomics as it provides direct protein evidence.

We reasoned that the integration of proteogenomics alongside Ribo-Seq could provide a solution to overcome the challenge incurred by ambiguous and multi-mapping reads during smORF identification during Ribo-Seq-only analysis. We refer to this as the Ribosome Profiling and Proteogenomics Pipeline (RP3). Although there are strategies to account for multi-mapping reads, such as ignoring them, splitting the reads across the mapped genes, or using statistical modeling of mapping uncertainty^16^, all of which are valid when working with sequencing reads alone, the integration of proteogenomics with Ribo-Seq is an advance since it provides evidence of translation by integrating mass spectrometry detection.

Using the RP3 workflow, we reanalyzed previously published datasets to provide additional evidence of smORFs that exist in regions inaccessible by Ribo-Seq alone (figure 1). Using the data from these results, we explore how Ribo-seq identifications differ from Liquid Chromatography with tandem Mass Spectrometry (LC-MS/MS) peptide evidence of novel translated microproteins, including differences in genome and protein sequence composition, transcript isoforms, gene paralogy, and the presence of repeat regions, all of which can preclude the identification of translational events. The data show that RP3 identifies smORFs and resultant microproteins that are disregarded in most pipelines that rely solely on Ribo-Seq evidence, significantly increasing the number Ribo-Seq detected smORFs with microprotein detection—the highest confidence set of microproteins.

**Figure 1.**
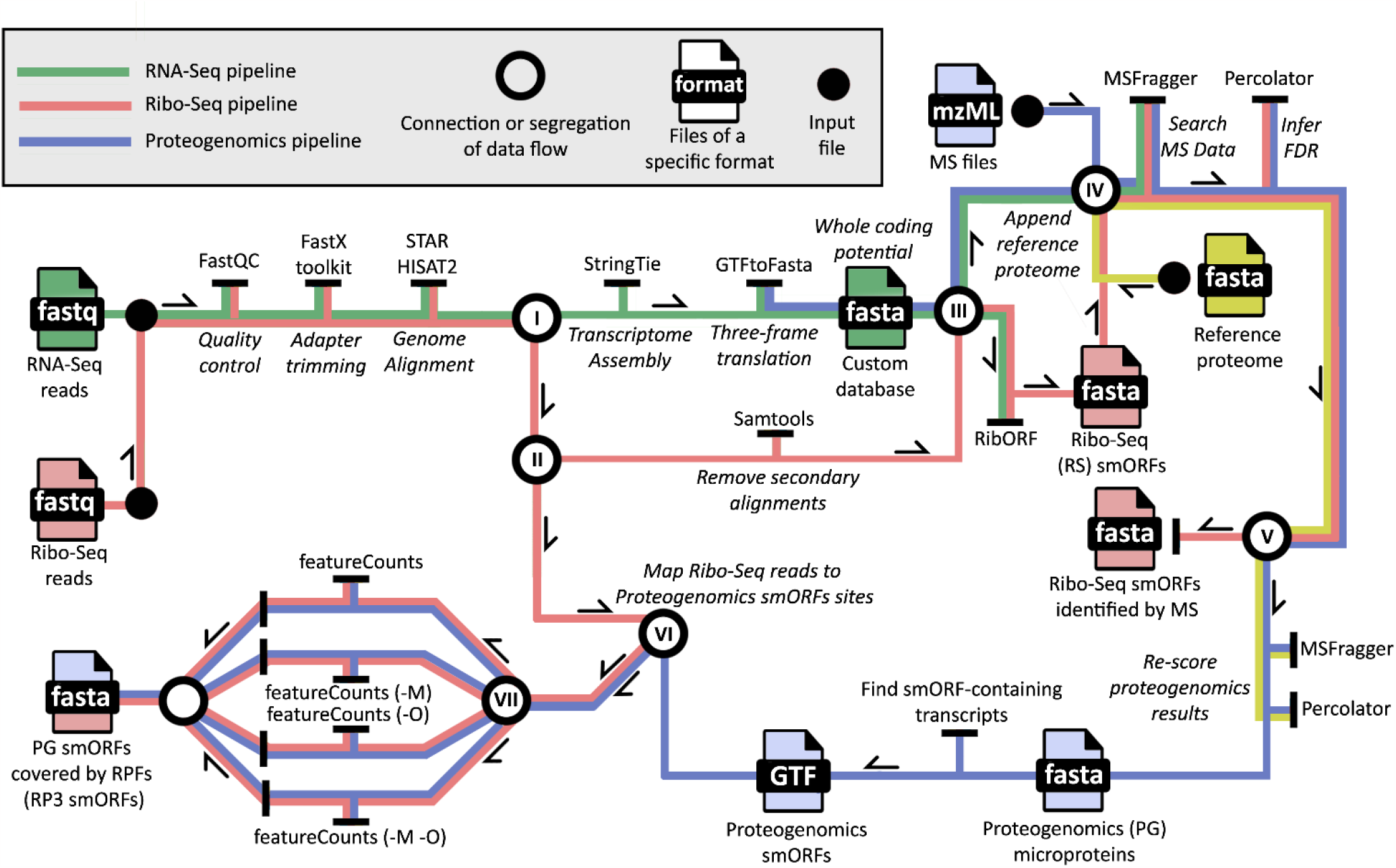
Workflow to reanalyze and overlay datasets. At the start of the workflow, RNA-Seq and Ribo-seq reads undergo the same initial steps, starting with quality control and adapter trimming, followed by the alignment to the genome with STAR^25^. Subsequently (I), the aligned RNA-Seq reads are used to assemble the transcriptome with StringTie^26^, and this assembly is translated to the three-reading frames to predict the whole coding potential of that transcriptome. (II) The aligned Ribo-seq reads are then divided into two subsets, one that contains all aligned reads, and another that contains only primary alignments, after removal of secondary alignments with samtools^27^. (III) The alignments filtered by samtools are scored against the three-frame translated database using RibORF, resulting in a fasta file containing Ribo-seq smORFs. A reference proteome is then appended to this fasta file to generate a custom database to check the mass spectrometry (MS) coverage for the Ribo-Seq smORFs. Similarly, the reference proteome is appended to the three-frame translated database, now without the Ribo-Seq smORFs, but with every predicted protein, which is the start of the proteogenomics pipeline. (IV) mzML files containing fragmentation spectra from LC-MS/MS experiments are searched against both databases using MSFragger^28^, whose results are filtered with Percolator^29^ to obtain an FDR of 1%. This results in two subsets of results (V), Ribo-Seq smORFs covered by both Ribo-seq and MS evidence, and proteogenomics-derived smORFs (PG smORFs), covered by MS evidence alone. The PG smORF-containing transcripts are located and (VI) the Ribo-seq reads containing secondary alignments are mapped to them using featureCounts^30^ (VII): once with default settings, to obtain PG smORFs covered by Ribo-seq reads, and then allowing ambiguous and multi-mapping reads to be included during read counting, resulting in another subset of PG smORFs covered by ambiguous and/or multi-mapped reads.

## Results

### Proteogenomics approach identifies microproteins that could not be found with Ribo-Seq

We have previously used Ribo-Seq to identify novel translated smORFs from thermogenic brown adipose tissue (BAT), energy storage-focused white adipose tissue (WAT), and beige adipose tissue^31^, which yielded a total of 3877 novel sequences, including a bioactive secreted microprotein encoded by the mouse gene Gm8773. In the study, the 3877 novel smORFs were translated and appended to the mouse UniProt database to create a custom proteomics database that is able to detect stable microproteins from these sequences. While these results validated the translation of 85 smORFs into microproteins, there is a vast difference in the numbers of microproteins obtained from Ribo-Seq and proteomics. Significantly higher numbers of smORFs from Ribo-Seq is common in the field, and typically only a handful of microproteins are detected using proteomics workflows that incorporate Ribo-Seq smORFs. The lack of detectable microproteins from Ribo-Seq annotated smORFs has been attributed to several factors including low abundance of microproteins, their length which results in the generation of few tryptic peptides, and lack of unique tryptic peptides. Indeed, we have detected microprotein via Western blot with antibodies that we have never obtained proteomics data from to further highlight some of the challenges of microprotein detection^32^.

Nevertheless, Ribo-Seq does not readily capture all ORFs. Alternative ORFs (AltORFs) overlap but are read in a different reading frame from the canonical ORF, and are easy to detect proteomically but difficult to detect by Ribo-Seq because of two different but overlapping reading frames that come from the same RNA. Several AltORFs have been shown to be functional^5^, highlighting the value in identifying these proteins. We also avoided the annotation of upstream overlapping smORFs (uoORFs) in the 3877 smORFs due to similar issues with overlapping transcripts, though uoORFs have been annotated in other RiboSeq studies^1,33^. To determine if we were missing any translated smORFs that were excluded from the Ribo-Seq dataset, such as AltORFs, we reanalyzed the data using a proteogenomics approach (Fig. 1a).

Proteogenomics relies on custom proteomics databases that are derived from the 3-frame translation of RNA-Seq data, which, in theory, should capture all translated smORFs from the translatome. After excluding peptides from the proteogenomics analysis that matched annotated proteins and keeping only the ones that matched predicted novel proteins exclusively, we identified 500 novel microproteins (Fig. 2a and Supplementary table 1). Although the number of Ribo-Seq identifications is eight times higher, the microproteins detected by proteomics are valuable since they are validated as stable members of the proteome. (Fig. 2b). After inferring the expression of the peptides in MS1 for both PG-derived and Ribo-Seq-derived (RS) microproteins, we found a significantly higher average intensity for PG microproteins (Fig. 2c).

**Figure 2.**
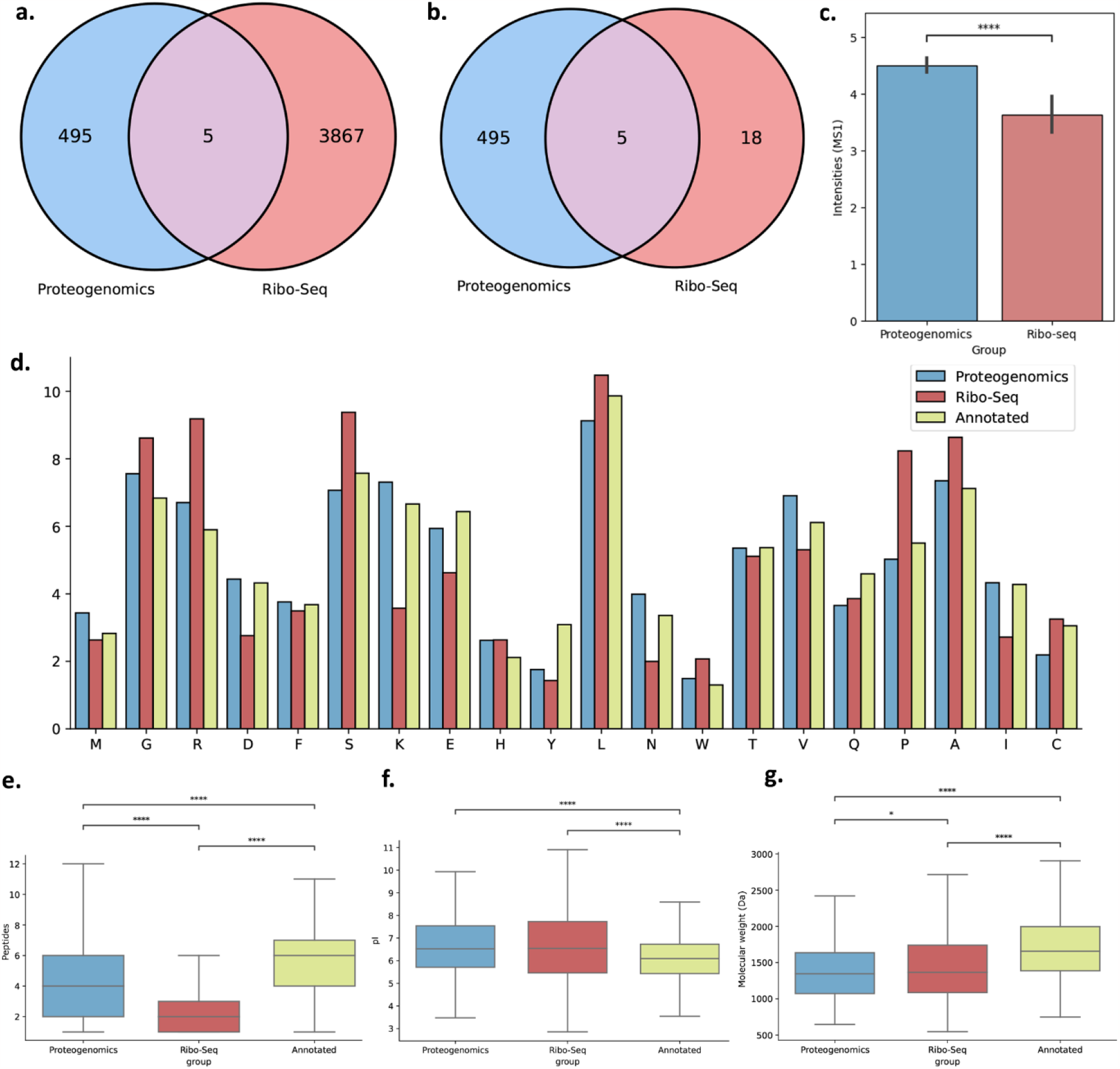
Comparison of microproteins identified by proteogenomics and Ribo-seq. Venn diagrams showing the number of novel microproteins identified by proteogenomics and Ribo-seq **(a)** and the intersection between proteogenomics microproteins and Ribo-seq microproteins that were confirmed by mass spectrometry **(b). c**, MS1 Ion intensities for peptides from proteogenomics and Ribo-seq microproteins (Mann-Whitney *\*\*\*P* = 1.53^-6^). **d**, amino acid distribution for each microprotein group. **e**, number of Unique Tryptic Peptides (UTPs) resulting from the *in silico* digestion of each microprotein using trypsin as the enzyme for each microprotein group (Kruskal-Wallis, *P* < 1.00^-4^. Dunn’s post-hoc *\*\*\*\*P* < 1.00^-4^). **f**, isoelectric points for each peptide resulting from the *in silico* digestion of each microprotein group (Kruskal-Wallis, *P* < 1.00^-4^. Dunn’s post-hoc *\*\*\*\*P* < 1.00^-4^). **g**, molecular weight for all three microprotein groups (Kruskal-Wallis, *P* < 1.00^-4^. Dunn’s post-hoc \**P* < 0.05, *\*\*\*\*P* < 1.00^-4^).

### Differences in sequence composition may confound mass spectrometry characterization of microproteins from Ribo-Seq results

Unannotated smORFs differ in their amino acid sequence composition when compared to already annotated proteins^2^. We wanted to see if this sequence bias extended to groups of microproteins coming from different approaches, such as proteogenomics and Ribo-Seq pipelines. Indeed, the distribution of amino acids varies among these groups (Fig. 2d), and lysine, which is of utmost importance for bottom-up proteomics^34^, is less common in the Ribo-Seq microproteins than it is for the ones found with proteogenomics, which could contribute to the difficulty in their identification by mass spectrometry. Proline, which is substantially enriched in Ribo-seq microproteins, is known to confound the interpretation of fragmentation spectra. For instance, if the residue is present at the C-terminal, it fragments almost exclusively by cleavage of the amide bond that is adjacent to that proline, resulting mostly in y1 fragment ions^35^.

Such amino acid biases could explain the low proteomics coverage for Ribo-seq microproteins, due to an increased difficulty in identifying microproteins rich in proline residues or those with very few tryptic sites. To confirm the latter, we inspected the unique tryptic peptides (UTPs) obtained after an *in silico* digestion of annotated, PG, and RS microproteins. The average number of UTPs is much larger for the annotated proteins, which makes them more likely to be found in a mass spectrometry experiment (Fig. 2e). The number of UTPs for proteogenomics microproteins is also higher than those coming from Ribo-seq, which could contribute to their higher proteomics coverage. The Isoelectric Point (pI) is also important for protein identification using Mass Spectrometry-based proteomics^34^. Similar to the number of UTPs, the pattern for these is also higher for PG microproteins when compared to the Ribo-seq (Fig. 2f). The pI for annotated microproteins, however, is significantly lower than the other two. Moreover, the molecular weight for all three microproteins groups also differs significantly (Fig. 2g).

### Most Ribo-Seq reads cannot be uniquely assigned to smORF-containing transcripts that encode the Proteogenomics-derived microproteins

To understand the minimal overlap between the proteogenomics and Ribo-Seq results we checked for evidence of translation for the proteogenomics hits. A priori we expected to find evidence for the translation of PG-microproteins but figured they were absent from the RS-microproteins because they fell below a threshold. We mapped Ribo-Seq reads for the PG-microproteins back to the transcripts that encode the smORFs. Surprisingly, we measured no Ribo-Seq coverage for the vast majority of proteogenomics smORFs when running FeatureCounts^30^ with the default settings (Fig. 3b), which should raise concerns about whether the peptide is a true identification or not, due to the lack of translational evidence, despite the presence of proteomics evidence. When we allowed the tool to include ambiguous and multi-mapping reads during read counting, however, we found translational evidence for many PG smORFs in the Ribo-Seq data.

**Figure 3.**
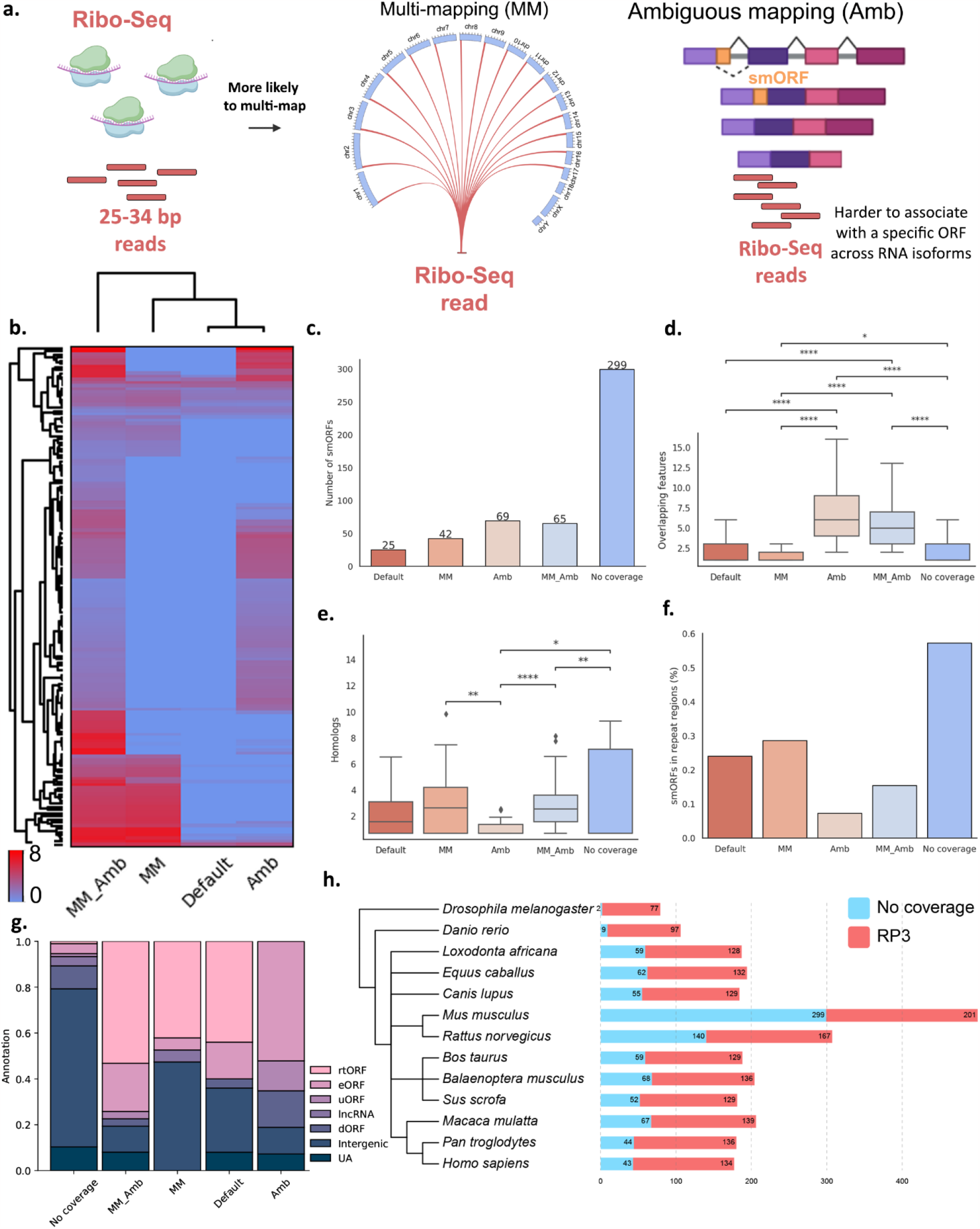
Ribo-Seq coverage for proteogenomics-derived (PG) smORFs. **a**, Schematic demonstrating the challenges imposed by shorter read lengths of Ribosome Protected Fragments (RPF) compared to RNA-Seq reads regarding ambiguous and multi-mapping. **b**, Heatmap showing the Ribo-Seq coverage (RPKM) for PG smORFs. Groups are classified based on the parameters allowed during read counting. Amb: featureCounts ran with the ambiguous mapping set to True `(-O)`; Default: featureCounts executed with default settings; MM: multi-mapping allowed `(-M)`; MM_Amb: both ambiguous and multi-mapping allowed `(-O -M).` smORFs that had higher expression in a specific cluster were grouped together for subsequent analyses. **c**, Absolute number of smORFs identified for each mapping group. Number of overlapping features (**d)** and homologs in the mouse genome (**e**) for the smORFs of each mapping group (Kruskal-Wallis *P* < 0.001, Dunn’s post-hoc *\*: P* < 0.05, *\*\*: P* < 0.01, *\*\*\*\*: P* < 1.00^-4^). No coverage includes smORFs with RPKM < 1 in all four mapping groups. **f**, Proportion of smORFs located in repeat regions for each mapping group. **g**, Annotation of smORFs for each mapping group based on the features they overlap in annotated transcripts. Intergenic: smORFs found within an intergenic region, i.e., smORFs that overlap no annotated gene. UA: Unknown Annotation. riORF: smORF located in a retained intron. dORF: smORF present downstream from an annotated ORF. uORF: smORF present upstream of an annotated ORF. lncORF: smORF present in a long non-coding RNA (lncRNA). rtORF: smORF present in a pseudogene or retrotransposon. eORF: smORF in an annotated exon that does not meet the previous criteria. **h**, Common NCBI tree for representative eukaryotes with bars to the side corresponding to the number of homologs in those species for the smORFs identified in this study with proteogenomics. Blue bars show conservation for PG smORFs with no Ribo-Seq coverage (RPKM < 1), and red bars show the number of conserved smORFs with Ribo-Seq coverage (RPKM > 1) when running featureCounts with either default settings, or with the parameters -O and -M turned on.

We grouped PG smORFs into clusters based on their detectability by Ribo-Seq. Some smORFs only have Ribo-Seq coverage when allowing for specific parameters during read counting, such as multi-mapping (Fig. 3a, middle), and ambiguous mapping (Fig. 3a, right) which means that these features preclude the identification of translational evidence for smORFs. Most smORFs do not have Ribo-Seq coverage when using default settings during read counting (Fig. 3b, 3c). We found a significant increase in the RPKMs for RPFs mapping to PG smORFs when allowing multi-mapping (MM), ambiguous mapping (Amb), or both (MM_Amb), and divided them into their respective mapping groups. smORFs and their respective mapping groups are described in Supplementary table 1.

We found that smORFs in both the Amb and MM_Amb groups are present in regions with a high number of transcript isoforms and/or other overlapping features (Fig. 3d), making it harder for the counting software to distinguish which transcripts the reads might be mapping to. To check whether genomic features were responsible for this, we inferred the number of paralogous sequences in the mouse genome for each smORF in these mapping groups. smORFs in the MM and MM_Amb group have a significant higher number of homologs in the genome compared to those in the Amb groups (Fig. 3e), which could keep RPFs reads from mapping to these regions due to a cutoff in the maximum number of mappings. Similarly, both the MM and MM_Amb groups also present a higher percentage of smORFs in repeat regions when compared to Amb (Fig. 3f). When not accounting for these features, the counting tool tends to disregard these reads, leading to an apparent lack of translational evidence. This data shows that PG smORFs, for which we have reliable protein evidence, would be disregarded by Ribo-Seq alone, due to limitations in the mapping step. By checking the coverage for PG smORFs while allowing for different mapping characteristics, we are able to detect translational evidence for these sequences as well. We refer to PG microproteins with Ribo-Seq coverage as RP3 microproteins. Those are backed by the strongest evidence, as we have both proteomics and Ribo-Seq coverage.

To better understand the genomic regions where these smORFs reside, we checked the annotated regions the smORFs overlap in their transcripts (Fig 3g). smORFs in the MM and Default clusters, as well as those that had no coverage, are located within intergenic regions most prevalently. smORFs affected by multi-mapping, in both the MM and MM_Amb clusters, commonly overlap the regions of pseudogenes and retrotransposons (rtORFs). Lastly, we checked the conservation levels for smORFs with no Ribo-Seq coverage and smORFs with coverage from the RP3 pipeline (Supplementary table 1). Overall, a higher proportion of RP3 smORFs are conserved across different eukaryotes (Fig. 3h), even in distant ones like *Danio rerio* and *Drosophilha melanogaster*, while the ones with no coverage have very few homologs in distant organisms. This suggests that the coverage inferred by our RP3 pipeline can identify the ones more likely to be evolutionarily conserved and thus more likely to be real.

### Mass spectrometry peptides allow the identification of smORFs with uncertain Ribo-Seq mapping

Using Ribo-Seq evidence alone would result in hundreds of microproteins being missing (Fig. 3b and 3c). By contrast, RP3 combines the strengths of Ribo-Seq and proteogenomics to identify a unique subset of smORFs with a very high level of confidence. As an example of the mapping landscape of short Ribo-Seq reads for PG smORFs, we plotted the position of each smORF identified with both proteogenomics and Ribosome profiling, along with the annotated genes using a circos plot (Fig. 4a). For instance, a smORF found on the reverse strand of chr9, (coordinates 44374744-44375091) would have been filtered out because of multi-mapping (Fig. 4b). The reads for this smORF mapped to more than 15 different regions in the genome and were filtered by the default parameters of STAR, which has a cutoff of < 10 secondary alignments, and by Ribo-Seq pipelines that use a cutoff of < 4 secondary alignments. These cutoffs are necessary for confident mapping due to shorter read lengths. By aligning the microprotein peptide sequence identified from a high-quality MS/MS spectrum (Fig. 4c) across the multi-mapped genomic locations (full multiple sequence alignment available in Supplementary figure 1) shows that the only nucleic acid sequence that is able to produce this particular microprotein is found at -chr9:44374744-44375091 (Fig. 4d). This example highlights the potential of RP3 to reveal microproteins with translational and proteomics evidence—the highest confidence group of smORFs/microproteins.

**Figure 4.**
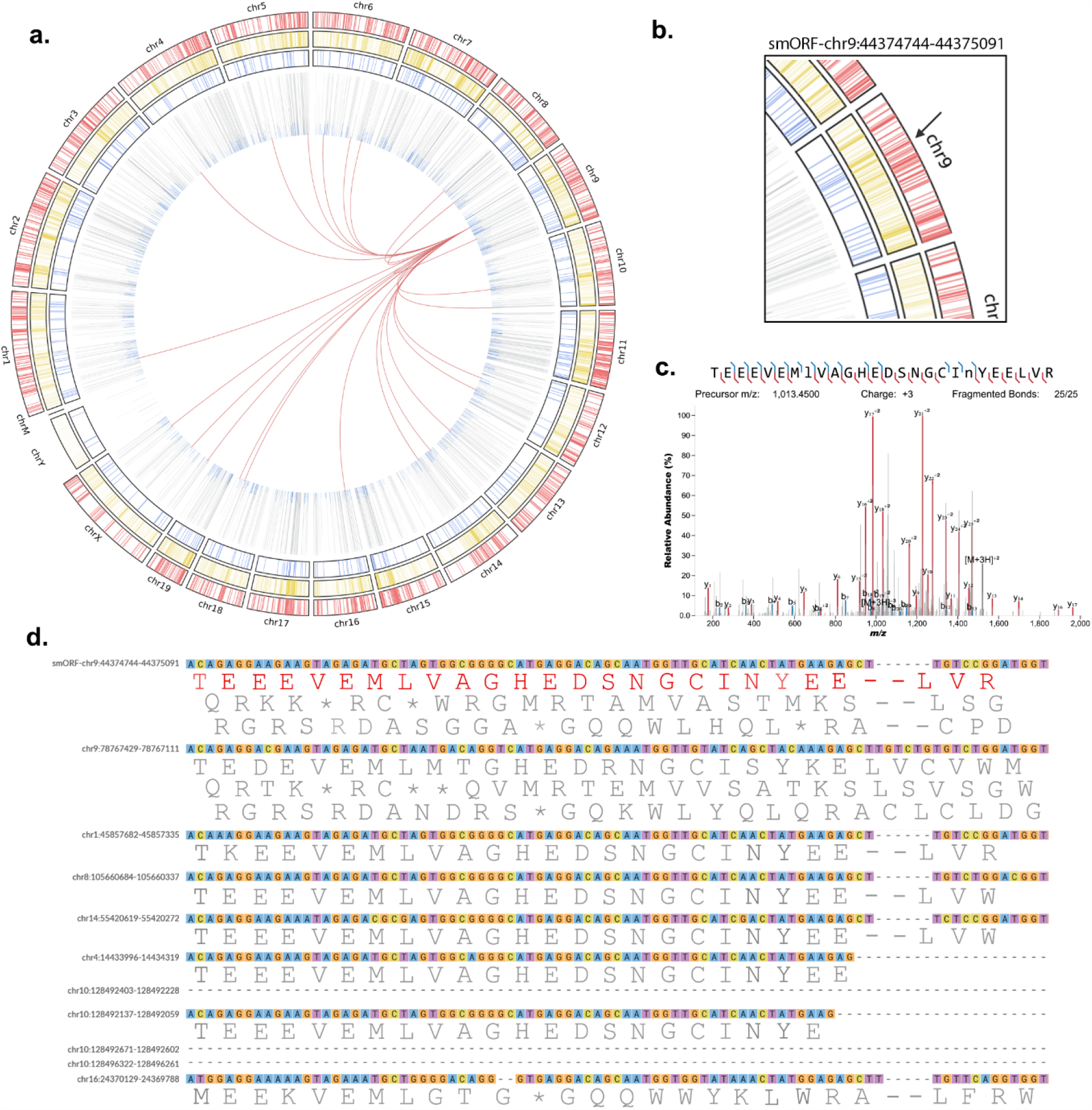
Mapping landscape of the representative smORF -chr9:44374744-44375091. **a**, Circos plot illustrating the genomic landscape of novel smORFs identified by Ribo-Seq (outermost ring, red bars) and proteogenomics (third ring from border to center, blue bars). The second ring from border to center contains the coordinates of annotated transcripts. The fourth ring contains peaks showing the total number of proteins a microprotein peptide maps to. Blue peaks indicate the presence of a tryptic peptide that does not match any predicted microprotein from the three-frame translation, thus representing truly unique-tryptic peptides (UTPs). The links in the center represent the other sites in the genome that the reads that map to the representative smORF also map to. **b**, Zoom-in of the representative smORF in the circos plot to highlight its genome position and features. **c**, Annotated fragmentation spectra showing the fragment b+ and y+ ions for a peptide of the microprotein encoded by the representative smORF. Lowercase letters represent amino acids with post-translational modifications. **d**, Multiple sequence alignment (MSA) of the nucleotide sequence of the representative smORF with all its homologs in the mouse genome. The peptide sequence from the fragmentation spectra in **c** is shown in its position in the nucleotide sequence of the smORF in the MSA.

### Proteogenomics allows the identification of a unique subset of peptides presented in the HLA complex

Peptidomics studies from human leukocyte antigen (HLA) complex provide a rich source of potentially immunogenic microproteins, and were used before to characterize smORFs identified by Ribo-Seq. Here, we reanalyzed published datasets^2,36^ to determine whether using RP3 would result in the identification of additional microproteins that were missed using the conventional pipeline (Fig. 5a). After translating the previously assembled transcriptomes to the three reading frames, we performed a proteogenomics search and identified 766 novel PG microproteins (Supplementary Table 2), 28 of which were shared between the proteogenomics and Ribo-Seq analyses (Fig. 5c). After checking for Ribo-Seq coverage using the same approach as in Fig. 2b, we found a similar pattern regarding mapping limitations (Fig. 5b), with hundreds of PG smORFs present in regions affected by ambiguous and/or multi-mapping (Fig. 5d).

**Figure 5.**
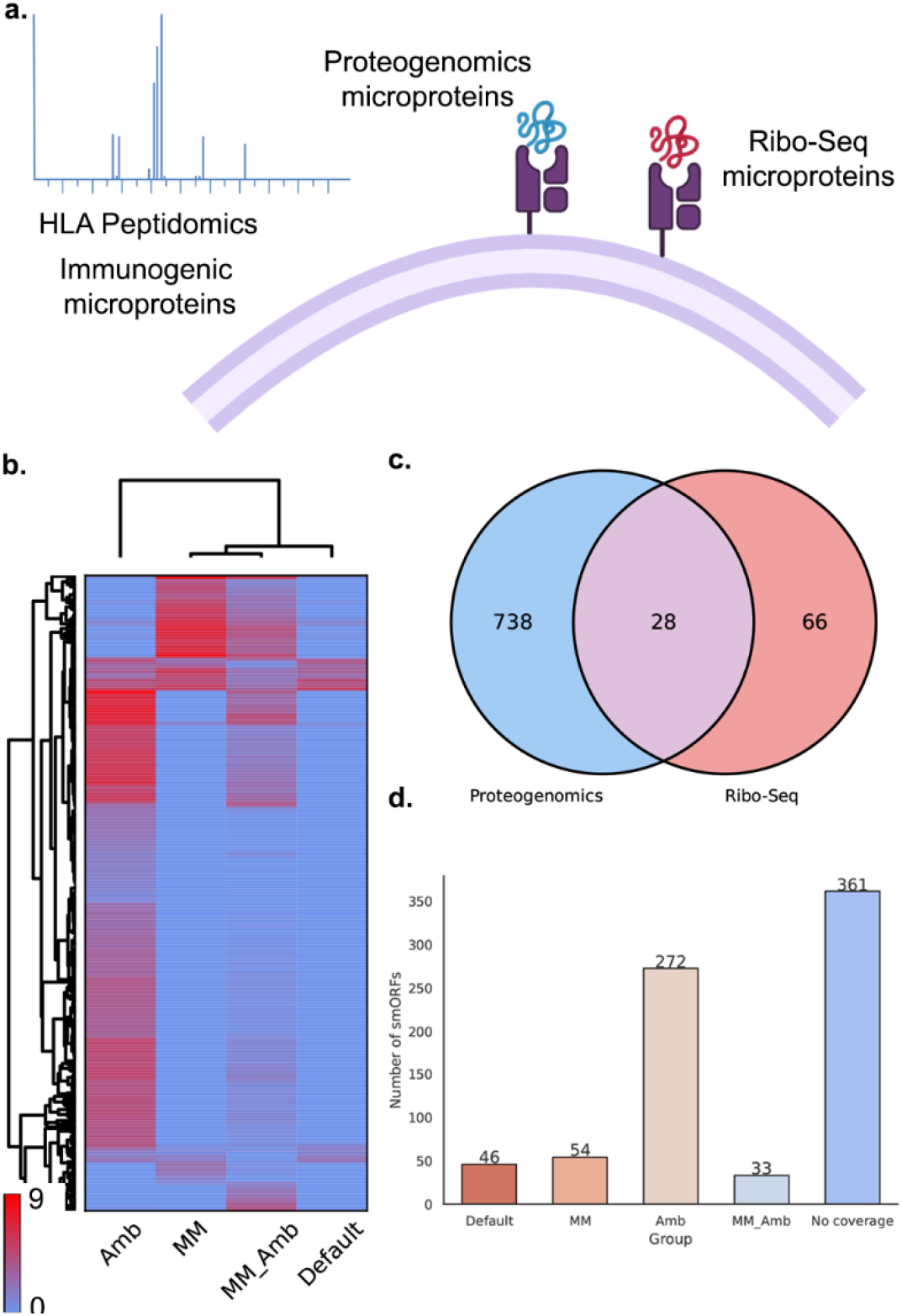
Identification of microproteins in HLA peptidomics datasets. **a**, Schematic drawing illustrating the presence of proteogenomics and Ribo-Seq microproteins presented on the HLA complex. **b**, Heatmap showing Ribo-Seq coverage for proteogenomics-derived (PG) smORFs. **c**, Venn diagram showing the intersection between PG and Ribo-Seq microproteins. **d**, Bar plot depicting the number of PG smORFs with Ribo-Seq coverage among the different clusters related to parameters allowed during read counting (same as Fig. 3b)

## Discussion

The orthogonal approach of using both proteogenomics and Ribo-Seq to identify smORFs we employed in this study is particularly important to perform a truly comprehensive microprotein identification analysis. We showed how limitations in the read alignment step of Ribo-Seq reads prevent the correct annotation of novel smORFs that can be found with proteogenomics unambiguously. Sequence-wise, there are two main reasons for this discrepancy: the presence of repeat regions and paralogous sequences for the smORFs in the genome, and multiple different mRNA isoforms that carry the same or a similar smORF.

Gene paralogy results in reads that map to multiple regions in the genome during the alignment step, called multi-mappers, which are often discarded after a certain threshold^16^. Reads that map to a genomic region that encodes many different transcript isoforms are often treated as ambiguous reads, which are disregarded by default by read counting tools such as featureCounts. Simply including all ambiguous and multi-mapping reads into the analysis is not advisable, however, as this would increase the number of false positives and add uncertainty about the coordinates the reads are in fact mapping to. Moreover, the number of uniquely assigned reads in our datasets is greatly undermined by the number of multi-mapping reads, which reinforces the idea of not simply including non-uniquely mapped reads. Proteogenomics solves that by allowing us to unambiguously assign a UTP to a microprotein in the custom database.

The annotated regions in the genome overlapped by novel smORFs provide some insights into the role of these novel sequences. In this study, instead of using traditional ways of annotating smORFs, we leveraged information that allowed us to better understand the reason behind the mapping limitations of Ribo-Seq very short reads. A big proportion of smORFs in MM and MM_Amb clusters were classified as rtORFs, meaning they share the loci with either pseudogenes or retrotransposons. This is very informative, as pseudogenes typically arise from the duplication of functional genes, and retrotransposons are present in multiple copies across the genome due to their highly dynamic copy-paste mechanism. Thus, these two classes of genes are expected to have high sequence similarity to other sites in the genome, resulting in regions prone to multi-mapping. eORFs are defined as those located in exons of already known transcripts and are prevalent in those regions with high numbers of isoforms. This is expected, since novel isoforms may give rise to exons that were previously deemed to not contain coding sequences. The low numbers of uORFs across all groups is especially informative, as this is the most common type of smORFs to be identified in Ribo-Seq studies. This means that proteogenomics and Ribo-Seq pipelines might be more suitable to identify distinct groups of smORFs each regarding their classification As proteogenomics hits are somewhat affected by false positives, and the number of PG smORFs with no Ribo-Seq coverage is considerably high in both proteomics datasets we analyzed, we advise the inclusion of PG microproteins that have at least coverage when allowing ambiguous and/or multi-mapping during read counting, to avoid the inclusion of hits that have no translational evidence whatsoever. PG microproteins with Ribo-Seq coverage (RP3 microproteins) and pure Ribo-Seq hits from established pipelines are thus considered the gold standard for microprotein identification in this study.

Moreover, to reduce the number of false positives, we performed a second round of searches during the proteogenomics analysis by generating a database consisting of annotated proteins plus the microproteins identified in the initial proteogenomics searches, which used the whole 3-frame translated database from the transcriptome. This way, the FDR assessment of proteogenomics hits is not that strongly affected by a bloated database consisting of millions of sequences and we can achieve an accurate global FDR. By reassessing the FDR and checking the Ribo-Seq coverage for PG microproteins, we are left with a very high-confidence set of novel sequences.

An important thing to note is that the overlap between the Ribo-Seq and proteogenomics results was negligible, as only five microproteins were shared between both approaches. This is surprising, as we expected a higher overlap between microproteins with proteomics evidence and microproteins with translational evidence. This reinforces the complementary aspect of these two different approaches, as each returns a unique subset of microproteins that could not be identified with the other method. Furthermore, the HLA peptidomics analysis provides a reliable source of evidence for novel microproteins, potentially enabling its annotation based on antigen presentation. Also, it shows that this pattern is neither limited to the mouse genome or to the presented Ribo-Seq and proteomics datasets from a specific study^31^.

## Conclusion

By integrating multi-omics datasets, we showed how proteogenomics can contribute to the identification of novel microproteins encoded by smORFs. Although Ribo-Seq is widely used for the study of the hidden proteome and is extremely powerful, it has some limitations in identifying microproteins derived from smORFs in regions where reads cannot uniquely map to, or that encode multiple RNA isoforms. After detecting multi-mapping reads, we presented a workflow to show how the Ribo-Seq coverage for a subset of smORFs is present, but most likely ignored by most bioinformatics pipelines. We also highlighted differences in sequence composition of microproteins identified with proteogenomics and Ribo-Seq, which could explain in part the difficulty of identifying a great overlap between the results of these two different approaches. We expect this workflow to increase proteomics coverage for Ribo-Seq results as well as to improve the quality of microprotein identification pipelines as those microproteins with Ribo-Seq and proteomics coverage are the most compelling for downstream biological studies.

## Methods

### Generation of custom databases for proteomics

To generate a database containing the whole coding potential of each cell line, we translated each transcriptome assembly generated by Martinez *et al*. (2023)^31^ to the three reading frames with the script GTFToFasta^2^, giving priority to ATGs. In the case of multiple ATGs for a region between two stop codons, the most upstream ATG was selected. Afterwards, we appended the Uniprot Mouse proteome (UP000000589) to the database and tagged each annotated sequence to be removed in the post-processing steps. For each resulting database, we generated a decoy database containing the reverse sequences of each protein plus the contaminants, which were appended to the initial database. We performed the same bioinformatic steps for the HLA peptidomics proteogenomics analysis, but used previously assembled human transcriptomes from HeLa-S3 and K562 cell lines^2^ to generate the custom databases, to which we appended the human reference proteome from Uniprot (UP000005640). For the mass spectrometry files, we downloaded raw data from the HLA peptidomics study from Bassani-Sternberg et al (2015)^36^, ProteomeXchange accession PXD000394.

### Peptide search and Post-processing of Mass Spectrometry data

We used MSFragger (v3.5) (ref. ^28^) to map fragmentation spectra to the custom database generated in the last step. The parameters were set as follows: precursor mass tolerance of 20 ppm, fragment mass tolerance of 50 ppm, Data-dependent acquisition as data type, fragment mass tolerance of 0.02 Da, and included carbamidomethylation (+57.021464 Da) as a fixed modification, and oxidation of methionine (+15.9949 Da) as a variable modification. For the validation step, we used Percolator (v3.06.1) (ref. ^29^) to infer the FDR with a cutoff set at 0.01 at the PSM and peptide levels. After the search, we removed any peptides that matched an annotated protein, but kept those that matched more than one predicted microprotein from the three-frame translated database. To make sure no annotated sequence was included in the results, we performed a Blastp (v2.12.0+) search against NCBI Refseq and excluded any microproteins that had a hit with identity and query cover 100%, i.e., a perfect match. Lastly, we performed a second round of searches to re-score the identified peptides by appending the final set of microproteins from the first search to the same reference proteome. Then, this smaller database was searched using the same mass spectrometry data and following the same bioinformatics steps. This was done to accurately assess the FDR without the effect of the bloated database coming from the three-frame translation of the transcriptome.

### Processing of Ribo-Seq data

We trimmed the Ribo-Seq reads to remove the Illumina adapters (AGATCGGAAGAGCACACGTCT) using fastx_clipper with the parameters `-Q 33, -l 20, -n, -v, -c`, piped to fastx_trimmed with the parameters `-Q 33, -f 1,` both tools from the FastX Toolkit (v0.0.13) (http://hannonlab.cshl.edu/fastx_toolkit/). Afterwards, we mapped the trimmed reads to the mouse mm18 or human hg19 genome to obtain a set of reads that did not map to the annotated regions. To do so, we ran the STAR aligner (v2.3.5a) (ref. ^25^) with `--outSAMstrandField intronMotif, -- outReadsUnmapped Fastx.` To remove reads mapping no tRNAs and rRNAs, we mapped the trimmed reads to a contaminant database running STAR with `-outSAMstrandField intronMotif, -- outReadsUnmapped Fastx`. Using as input the unmapped reads in fastq format, we ran STAR with the parameters `--outSAMstrandField intronMotif, -outFilterMismatchNmax 2, -- outFilterMultimapNmax 99, --chimScoreSeparation 10, --chimScoreMin 20, -- chimSegmentMin 15, --outSAMattributes All.` We specified a max of 99 multi-mappings so STAR would not discard all reads mapping up to that many regions, as it does not keep reads with a number of alignments above the specified threshold.

### Analysis of ambiguous and multi-mapping reads

To check for Ribo-Seq coverage for the smORFs we could find peptide evidence for with proteogenomics, we first appended the coordinates of those smORFs to the mouse mm18 reference `GTF` file. Then, we performed four rounds of read counting with featureCounts (v1.6.3) (ref.^30^), first with default parameters and then allowing multi-mapping reads with `-M,` allowing ambiguous mapping reads with `-O,` and allowing both ambiguous and multi-mapping reads with `-M` and `-O`. We plotted the counts for each combination of these two parameters (-M: MM, -O: Amb, -M -O: MM_Amb, and Default, when using default parameters) into a heatmap using the Python package nheatmap (https://pypi.org/project/nheatmap/). We selected the smORFs whose read counts were higher in a specific setting and grouped them together, except if the smORF had a RPKM > 1 using default settings. In that case, the smORF was classified as “Default”, regardless of its coverage when allowing the other parameters. For each group, we performed comparisons regarding homology, number of isoforms, and repeat regions as follows: first, to infer homology, we performed a blastn search using each smORF as query and the genome as subject, selecting only alignments with an e-value < 0.001 and a score > 50, and generated box plots accordingly. For the overlapping features, we ran bedtools intersect (v2.30.0) (ref. 31) using the reference GTF file to identify how many features overlapped each smORF, and plotted those into box plots. To investigate repeat regions, we used repeat files for the mm10 genome provided by RepeatMasker (http://www.repeatmasker.org/species/mm.html) and checked which smORFs overlapped the repeat coordinates. For overlapping regions, we used a custom script to classify the overlap with the reference GTF file for mm10 from Ensembl after running bedtools intersect on the reference and custom GTF file containing the smORFs. To analyze the multi-mapping landscape, we performed additional filtering in the `sam` files to analyze the distribution of reads with secondary alignments across the genome. For each `sam` file, we first kept only the reads with the flag 0×100, which indicates a secondary alignment, by running Samtools (v1.13) with the parameter `-f 0×100`. Afterwards, we used a custom script, `mm_inspector.py`, to iterate the alignments and select the reads that aligned at least once to a region in the genome that encodes a proteogenomics smORF based on the coordinates of our custom `GTF` file, while keeping all the other regions these selected reads mapped to.

### Circos plot generation

To generate the circos plot for read mapping visualization at the genome level, we used the Python package PyCircos (v0.3.0) (https://github.com/ponnhide/pyCircos) and added bars corresponding to the coordinates of each gene annotated in the reference mm10 `GTF` file to the outermost ring. Then, we did the same for the smORFs found with Ribo-Seq and for the Proteogenomics smORFs, which were added to the second and third ring, from edge to center, respectively. Using a filtered sam file containing only reads that mapped to at least one Proteogenomics smORF, we selected the reads that mapped to the representative smORF. Afterwards, we added links to the center of the plot whose coordinates they point to in the genome correspond to the regions the reads map to. To generate the multiple sequence alignment, we first identified homologs by running tblastn on the mouse genome assembly mm10 and selected hits with an E-value < 0.001 and a bitScore > 50. Then, we performed the multiple sequence alignment with these sequences using MAFFT^37^.

### Sequence analysis comparison among annotated, Ribo-Seq and proteogenomics microproteins

To plot the amino acid composition, we extracted the microproteins sequences for each group of microproteins and plotted the distribution with the Python package matplotlib. To obtain the number of UTPs for each microprotein and the isoelectric point for each peptide, we used the tool RapidPeptidesGenerator (RPG)^38^, using trypsin as the enzyme for the *in silico* protein digestion, and plotted the distributions with matplotlib.

### Conservation analysis

To identify PG smORF homologs in the genome of other eukaryotic organisms and check their overall level of conservation, we a performed a tBlastn search against the genome assemblies of *Equus caballus* (GCF_002863925.1), *Homo sapiens* (GCF_000001405.40), *Canis lupus (*GCF_000002285.5), *Sus scrofa* (GCF_000003025.6), *Bos taurus* (GCF_002263795.2), *Pan troglodytes* (GCF_028858775.1), *Rattus norvegicus* (GCF_015227675.2), *Danio rerio* (GCF_000002035.6), *Balaenoptera musculus* (GCF_009873245.2), *Loxodonta africana* (GCF_000001905.1), *Drosophila melanogaster* (GCF_000001215.4), *Macaca mulatta (*GCF_003339765.1). Then, we filtered the alignments to include only those with an E-value < 0.001 and a bitScore > 50. We then annotated a common NCBI tree in phylip format for these species using EvolView^39^ by adding bars to the side of the tree corresponding to the numbers of smORF homologs in the organism, according to their Ribo-Seq coverage.

### Statistical analysis

We used the Python package SciPy (https://scipy.org/) to perform statistical analysis when adequate. We first checked for a Gaussian distribution for each group using the Shapiro-Wilk test. Since our samples followed a non-parametric distribution, we ran the Kruskal-Wallis test followed by Dunn’s post hoc adjusted for multiple hypothesis testing with the Benjamini-Holchberg FDR correction using SciPy, and added statistical significance to the plots using statsannotations (https://github.com/trevismd/statannotations).

## Data availability

Raw data from Ribo-Seq and mass spectrometry experiments were previously generated by Martinez et al. (2022) ^25^ and downloaded from Gene Expression Omnibus (GEO Series: GSE198109) and MassIVE (MSV000089022), respectively. smORF sequences previously identified by Ribo-Seq and raw Ribo-Seq data were obtained from Martinez et al (2020)^2^ (GEO accession <u>GSE125218</u>). HLA peptidomics datasets were previously generated by Bassani-Sternberg (2015)(Ref. ^36^) and downloaded from proteomeXchange (accession PXD000394). Information regarding novel microproteins identified in this study is available in Supplementary Tables 1 and 2.

## Acknowledgements

A. Saghatelian acknowledges financial support from NIH grants <u>P30CA014195</u>, <u>R01GM102491</u>, and <u>RC2DK129961</u>; Frederick Paulsen and the Ferring Foundation; and a sponsored research agreement with Novo Nordisk Research Center Seattle, Inc. This work was supported in part by National Institute of Science and Technology on Tuberculosis (Decit/SCTIE/MS-MCT-CNPq-FNDTC-CAPES-FAPERGS) [grant number 421703/2017-2], Banco Nacional de Desenvolvimento Econômico e Social (BNDES/FUNTEC) [grant number 14.2.0914.1], FAPERGS [grant numbers 17/1265-8 INCT-TB and 19/1724-3 PQG] and CAPES (CAPES-Print program 041/2017). C.V.B (CNPq, grant 311949/2019-3) is a Research Career Awardee of CNPq. This study was financed in part by the Coordenação de Aperfeiçoamento de Pessoal de Nível Superior—Brasil (CAPES), Finance Code 001. We would like to acknowledge the financial support given by Coordenação de Aperfeiçoamento de Pessoal de Nível Superior—Brasil (CAPES). CVB would like to acknowledge financial support given by CNPq/FAPERGS/CAPES/BNDES to the National Institute of Science and Technology on Tuberculosis (INCT-TB), Brazil. Templates from Biorender.com were used to generate Figures 3 and 5.

## Author contributions

E.V.S and A.S conceptualized the study. E.V.S., C.A.B., B.M. and A.L.B. performed the bioinformatics methodologies, visualization and data curation. E.V.S. wrote the python code for the bioinformatics pipeline. A.S., C.V.B, P.M., and L.A.B. performed supervision of the research. E.V.S., A.S., and C.V.B. wrote the original draft of the manuscript.

## Supplementary information

Supplementary Table 1: Microproteins identified by mass spectrometry and covered by Ribo-Seq reads from mouse adipose tissue.

Supplementary Table 2: Microproteins identified by mass spectrometry and covered by Ribo-Seq reads from HLA peptidomics datasets.

Supplementary figure 1: Full multiple-sequence alignment for the representative smORF and its homologs in the mouse genome.

