## Supplementary figures and images for "The Integration of Proteogenomics and Ribosome Profiling Circumvents Key Limitations to Increase the Coverage and Confidence of Novel Microproteins"

### Supplementary figure 1

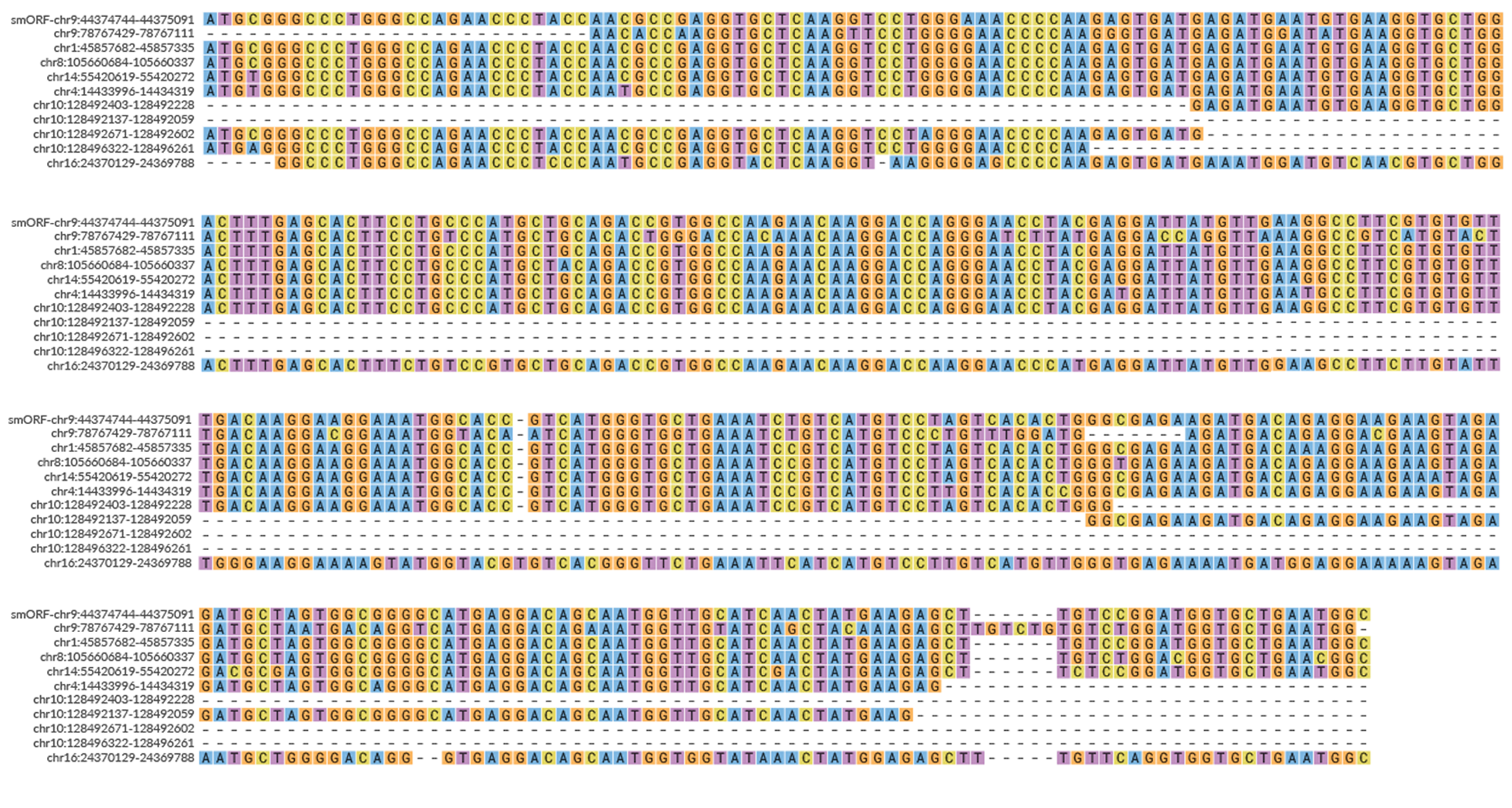
